# Beetle bioluminescence outshines aerial predators

**DOI:** 10.1101/2021.11.22.469605

**Authors:** Gareth S. Powell, Natalie A. Saxton, Yelena M. Pacheco, Kathrin F. Stanger-Hall, Gavin J. Martin, Dominik Kusy, Luiz Felipe Lima Da Silveira, Ladislav Bocak, Marc A. Branham, Seth M. Bybee

## Abstract

Bioluminescence is found across the tree of life and has many functions. Yet we understand very little about its timing and origins, particularly as a predator avoidance strategy. Understanding the timing between bioluminescence and predator origins has yet to be examined and can help elucidate the evolution of this ecologically important aposematic signal. Using the most prevalent bioluminescent group, fireflies, where bioluminescence primarily functions as aposematic and sexual signals, the timing for the origins of both potential predators of fireflies and bioluminescence is explored. Divergence time estimations were performed using genomic-scale phylogenetic reconstructions, and multiple fossil calibration points, allowing for a robust estimate for the origin of lampyrid bioluminescence as a terrestrial and as an aerial signal. Our results recover the origins of terrestrial beetle bioluminescence at 141.17 (122.63–161.17) mya and firefly aerial bioluminescence at 133.18 (117.86–152.47) mya with a large dataset focused on Lampyridae; and terrestrial bioluminescence as 148.03 (130.12–166.80) mya, with the age of aerial bioluminescence at 104.97 (99.00–120.90) mya with a complementary broad Elateroidea dataset. These ages predate the origins of all known extant aerial predators (i.e., bats and birds) and support the much older terrestrial predators (assassin bugs, frogs, ground beetles, lizards, snakes, hunting spiders, and harvestmen) as the most likely drivers of terrestrial bioluminescence in beetles, and sexual signaling likely being the original function in aerial fireflies.

## Introduction

Bioluminescence has evolved independently almost 100 times across both eukaryotes and prokaryotes (e.g., insects, crustaceans, other marine invertebrates, fish, protists, fungi, and bacteria; [1, 2, 3,4, 5, 6]). The function of bioluminescence varies across organisms including: prey attraction, [7, 8, 9, 10] predator avoidance, [11, 12], counterillumination [13, 14], sexual communication [15, 16], and spore dispersal [17, 18]. In the terrestrial environment, bioluminescence is principally used as an aposematic signal [11, 19, 20, 21, 12, 22].

While Lampyridae (fireflies) are perhaps the most well-known group of bioluminescent beetles, there are four additional extant families that contain bioluminescent species in the superfamily Elateroidea: Elateridae, Sinopyrophoridae, Phengodidae, and Rhagophthalmidae. Two independent origins of bioluminescence in Elateroidea were suggested by the phylogenetic analysis of Martin et al. [23] and Kusy et al. [24]. One origin of bioluminescence in the common ancestor of fireflies and their relatives (Lampyridae, Rhagophtalmidae, Phengodidae and Sinopyrophoridae) was supported by Kusy et al. (24), and a second independent origin in click beetles (Elateridae). Evidence for these two origins is corroborated by Fallon, Lower et al. [25] who provided genomic evidence that strongly supports the independent evolution of elaterid and lampyrid luciferases. We focus our analysis on the origin of bioluminescence in the lampyroid group, including the families Lampyridae, Phengodidae, and Rhagophthalmidae. All known lampyroid larvae are bioluminescent [26], suggesting larval bioluminescence is the ancestral state for each of these families and their common ancestor. These larvae are also largely terrestrial [15].

Current evidence suggests that larval elateroid bioluminescence likely originated as a predator avoidance strategy in fireflies [23, 27, 28, 29, 30, 31] by allowing these ground-dwelling invertebrates to advertise their chemical defenses to potential predators [12, 20, 21, 29, 31]. For example, Underwood et al. [32] found that firefly predators learned to avoid distasteful prey when it was associated with light, ultimately resulting in effective predator avoidance. This demonstrates the effectiveness of bioluminescence, when accompanied by chemical defenses, as a strong aposematic signal.

Independent from larval bioluminescence, adult firefly bioluminescence (with light organs in different abdominal segments: [27]) subsequently evolved separately in several firefly lineages [23, 27, 33]. Today, this adult bioluminescence is mainly used for sexual communication in the form of complex aerial bioluminescent courtship displays by male fireflies, followed by a response from usually sedentary conspecific females on the ground or in the vegetation above [34]. There is evidence that adult firefly bioluminescence also accelerates avoidance learning by potential predators and thus also functions as an aposematic signal for aerial predators (*i*.*e*. bats; [35, 36]). However, it is unclear how adult firefly bioluminescence originated. In fact, despite the modern interactions between larval and adult fireflies with many different predator groups [37, 38], and a few claims that specific predators caused adult firefly bioluminescence [36], none of these groups can yet be attributed to the origin of bioluminescence as an aposematic signal in fireflies.

To investigate the origins of bioluminescence in the predator context, we distinguish between terrestrial bioluminescence and aerial bioluminescence to test which predator groups may have driven the origin of larval bioluminescence in the common ancestor of Lampyridae (fireflies), Rhagophthalmidae (railroad worms), and Phengodidae (glowworms), and which predator groups may have contributed to the origin of adult bioluminescence in fireflies. From a predator perspective, adult larviform or wingless (apterous or brachypterous) females tend to be active on the ground and thus display terrestrial bioluminescence. The origin and evolution of terrestrial bioluminescence as an aposematic signal would have likely been driven by terrestrial predators (e.g., carabid beetles, arachnids, centipedes, amphibians, reptiles, and rodents; [37]. In contrast, winged (pterous) adult males use aerial bioluminescence (above ground) in their complex and highly visible courtship displays, and their winged adult females also tend to respond with individual flashes from higher up in the vegetation (above ground or aerial bioluminescence). For aerial bioluminescence to arise as an aposematic signal, aerial predators (e.g., bats, dusk/night active birds; [39, 37]) would have to pre-date or coincide with the origin of aerial bioluminescence displays. Most importantly, to contribute to the origin of terrestrial and/or aerial (above-ground) bioluminescence in fireflies, a potential predator group would have to pre-date or coincide with the respective origins of terrestrial or aerial bioluminescence.

Here we estimate when terrestrial, larval beetle bioluminescence (Lampyridae, Phengodidae, Rhagophthalmidae) originated, as well as the time of the first origin of adult, aerial bioluminescence in fireflies (Lampyridae). We take advantage of eight described fossils and two genomic-scale phylogenies [40, 41] to date the origins of bioluminescence in elateroid beetles. Both phylogenies were focused on elateroid beetles, but with a different emphasis in taxon sampling. Martin et al. [40] focused on subfamilies and tribes within Lampyridae while Douglas et al. [41] focused on subfamily relationships within the Elateridae. By dating each of these two topologies independently, we can take into account how taxon sampling and fossil placement may affect the resulting divergence time estimates, thereby providing the most robust estimate for the age of bioluminescence in this group to date. We use the resulting age estimates to examine the presence of potential terrestrial and aerial predator groups at the estimated origins of terrestrial bioluminescence in beetles and aerial bioluminescence in fireflies to test hypotheses on the roles of these predators in the origin of bioluminescence. These results shed light on potential selective agents for the origin of beetle bioluminescence, both on the ground and in the air, ultimately giving rise to some of the most diverse and captivating light displays seen in the terrestrial environment.

## Material and methods

### Phylogenies used for divergence time estimation

Bayesian divergence time estimation combines phylogenetic hypotheses with prior knowledge of molecular clocks or, in this case, the fossil record allowing for investigations into the time of origin for organisms and their ecological innovations [42, 43]. Fossil calibrations are known to have a significant impact on divergence time estimation depending on their placement and the ages provided by the user [44, 45]. We obtained both the tree file and alignments from Martin et al. [40] and the alignments provided by Douglas et al. [41] to perform divergence time estimation. The alignments from Douglas et al. [41] were ran in ModelFinder [46] and then used to reconstruct a topology in IQ-Tree v1.6.12 [47] utilizing ultrafast bootstraps [48] and compared with the published phylogeny to confirm congruence.

We used *chronoPL* as implemented in the R package [49] APE [50] to produce fixed topologies of the Martin and Douglas trees for divergence time analyses [51, 52]. Using a fixed topology cuts down on the necessary computing time and ensures that the resulting ages are not impacted by minor differences in topology. This program allowed us to transform the topologies into ultrametric trees and adjust the node heights such that they were within the bounds of the fossil prior distributions we provided in subsequent divergence time estimation (see *Fossil selection and placement* below).

### Fossil selection and placement

We selected a total of nine fossils with a focus on those that could provide reliable calibration points for major clades across each topology. Two of these fossils were used on both topologies. The placement of the fossils was determined based on morphological similarities with extant taxa and currently proposed placements (see Supp. Table 1). Other described fossils were not included (e.g., [53, 54]) when older fossil representatives in their corresponding clades were available. Relative ages for each fossil were obtained by the Paleobiology Database [55], which uses stratigraphic information to assign ages to geologic deposits.

For the Martin et al. [40] topology, four fossil priors were placed based on the published classification and study of the original descriptions and images. One was placed at the base of Luciolinae (*Protoluciola albertalleni*; [56]) and one at the node Phengodidae + Rhagophthalmidae (*Cretophengodes azari;* [57]), a third fossil was placed at the base of the subfamily Lamprohizinae (*Phausis fossilis*; [58, 59]), and the fourth at the base of the *Photinus* (*Photinus kazantsevi*; [60]). Due the questionable assignment of *P. kazantsevi* to the genus *Photinus* and its therefore problematic placement within the phylogeny (Branham and Silveira unpublished data), we also ran a subset of analyses on the Martin topology excluding this fossil. Removal of the *Phontinus* fossil prior had minimal impact on the ages for surrounding nodes (−1 to

+8 ma). For the Douglas et al. [41] topology, we used two fossil priors that overlapped with the calibration points used for Martin et al. [40]. These were placed at the base of Lampyridae (*Protoluciola albertalleni*; [56]) due to a lack of species determinations used by Douglas et al. [41]. The other was once again placed at the base of Phengodidae + Rhagophthalmidae (*Cretophengodes azari;* [57]). Additionally, we placed four fossils to constrain major Elateridae clades across the topology: one at the base of the Agrypninae (*Ageratus delicatus*; [61]), one at the base of the Lissomini (*Lissomus taxodii*; [61]), one at the base of the Negastrini (*Ganestrius elongatus*; [61]), and one at the base of *Cardiophorus* (*Cardiophorus exhumatus*; [61]). Each of these four fossils was assigned a soft maximum of 242 my based on the oldest Elateridae (*Elateridium* spp.; [61]).

When using fossil specimens as calibration points, we used an exponential probability distribution as it allows relative ages of each fossil to be used as “hard” minimums, as there is no need to sample ages younger than the fossil evidence being used. In contrast, maximum ages are “soft” allowing the divergence time estimation to sample ages older than that proposed maximum, but do so by decreasing the probability of those ages as they get further from that maximum age [62]. Furthermore, an exponential distribution is preferred in the absence of additional information as it requires only two parameters (minimum age and the mean) to be set by the user over the three required for a log normal distribution [62]. We generated potential age distributions by setting the relative age of the fossil as the hard minimum, and adjusting the mean age such that the 95% quantile of the exponential distribution matches that of our soft maximum, but does not exceed the age of the oldest known representative of the parent clade (i.e., the maximum age for the subfamily Agrypninae does not exceed the age of the oldest known representative for the family Elateridae).

### Divergence time estimation

Next, we employed BEAUTi [63] to set all analysis parameters such as fossil placement and age distributions, fixed starting trees, and the tree and clock models. To account for sensitivity to model choice, four different analyses were performed on each dataset (i.e., Martin and Douglas). Each analysis used a different combination of tree and clock models which included: Birth-Death (BD) and Relaxed Clock Exponential (RCE), BD and Relaxed Clock Log Normal (RCLN), Yule and RCE, Yule and RCLN. Analyses were performed. Divergence time estimation, utilizing the files prepared in BEAUTi, were run in BEAST v.2.6.0 [63] via the CIPRES Science Gateway v.3.3 (www.phylo.org). To ensure convergence of our Bayesian analyses, and to determine burn-in, the resulting log file for each analysis was viewed in Tracer v.1.7.1 [64]. Analyses were run until Effective Sample Sizes (ESS) of parameters of interest were >100, with many parameters being >200 (this required a chain length of 200,000,000 and 450,000,000 for Martin and Douglas respectively). Lastly, TreeAnnotator v.1.10.4 [42] was used to generate consensus ages for each analysis after a burn-in (10–50%) was discarded. All ages are reported as the median age followed by the 95% highest posterior density interval (HPD).

### Ancestral state reconstruction

We evaluated the origin of bioluminescence for both the larval and adult life stages. Larval bioluminescence was coded as absent (0) or present (1). In these analyses larval bioluminescence was considered equivalent to adult terrestrial bioluminescence because all known larval fireflies lack wings and thus would not be expected to be displaying in the aerial environment during this life stage (Lloyd, 1983). Adult bioluminescence was coded as either absent (0), female-only terrestrial bioluminescence, with non-bioluminescent males (1), or aerial bioluminescence of either males and females (2) (Supp. Table 2). Maximum parsimony and maximum likelihood ancestral state reconstructions were conducted for both data sets with Mesquite v3.61 [65].

### Potential predators

If larval beetle bioluminescence arose to advertise a chemical defense in terrestrial larvae then there must have been predation pressure by terrestrial predators selecting for this trait. If adult firefly bioluminescence arose as an aposematic signal during aerial displays then there must have been aerial predators preying upon these aerial bioluminescent individuals. Terrestrial and aerial animal groups known to prey on fireflies (e.g. insectivores) were compiled from [37, 66], and authors’ personal observations (see results). Extinct insectivorous groups (e.g., Pterosauria) were also considered. To assess the potential role of each predatory group in the origins of terrestrial (larval and flightless females) and aerial adult beetle bioluminescence, predator group ages were compared with our estimates for the origin of terrestrial and aerial bioluminescence. The ages of each firefly predator group were gathered from the literature [45, 67, 68, 69, 70, 71, 72, 73, 74] (Supp. Table 3).

## Results and Discussion

Terrestrial (larval) bioluminescence is an ancestral trait in Lampyridae (and their relatives: Phengodidae, and Rhagophthalmidae), preceding the origin of adult aerial bioluminescence (Figure 1) (see also Supp. Figures 1, 2, 3, and 4). Our results recover the origin of terrestrial beetle bioluminescence at 141.17 (122.63–161.17) mya. This was followed by the origin of aerial bioluminescence in fireflies (Lampyridae) at 133.18 (117.86–152.47) mya (Figure 1). These ages are corroborated by our analysis of the Douglas et al. [41] dataset. These independent divergence time estimates recovered the origins of terrestrial bioluminescence at 148.03 (130.12–166.80) and aerial bioluminescence at 104.97 (99.00–120.90). Both sets of recovered ages are older than previous estimations and preclude modern aerial predators as selective agents for the origin of beetle bioluminescence.

**Figure 1.**
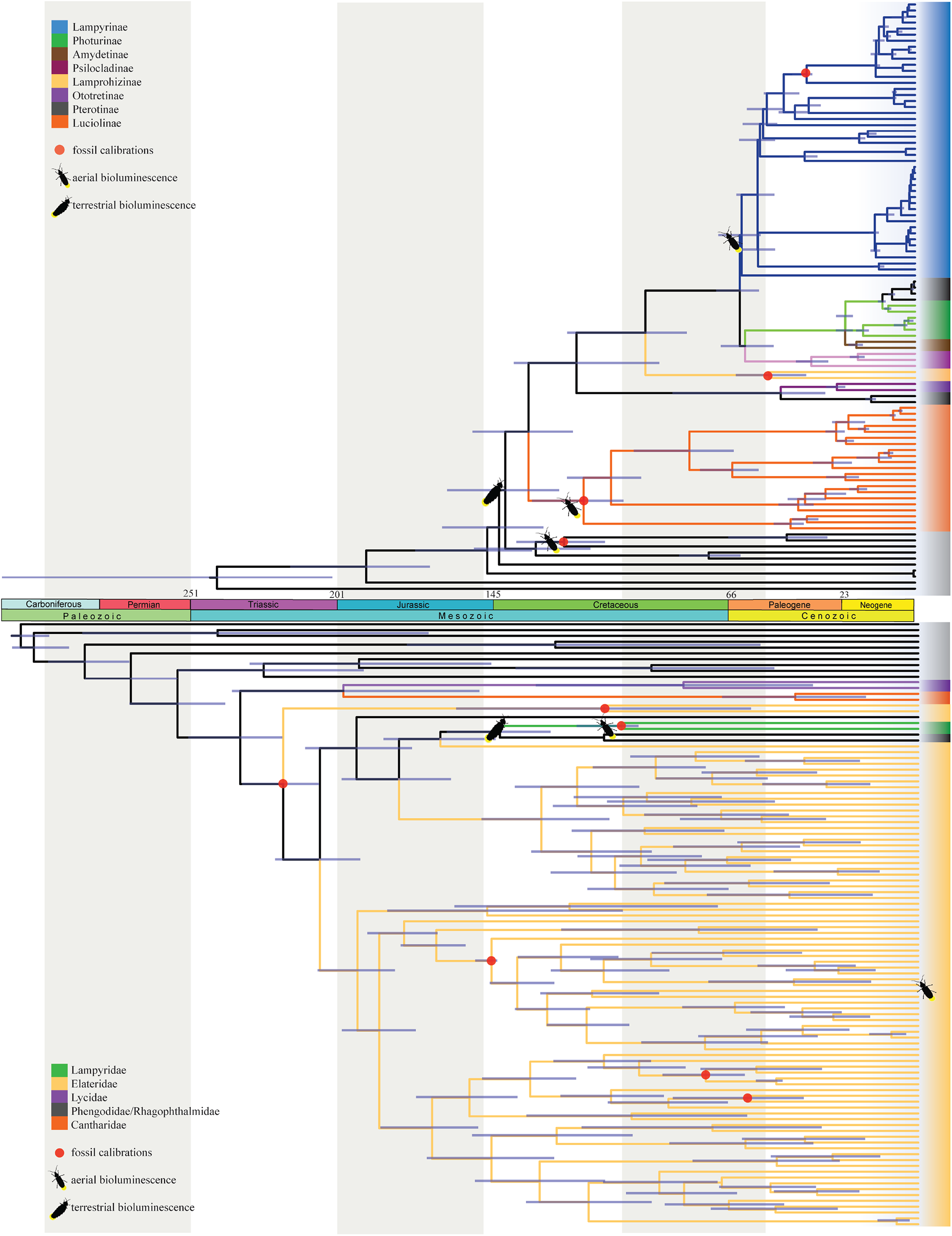
Two divergence-time calibrated phylogenies of Elateroidea using LN and BD (based on data from Martin et al., [40], top, and Douglas et al., [41]. bottom) with results of ancestral state reconstruction for both terrestrial and aerial bioluminescence depicted at appropriate nodes. Gray vertical bars represent 50 million years.

**Figure 2.**
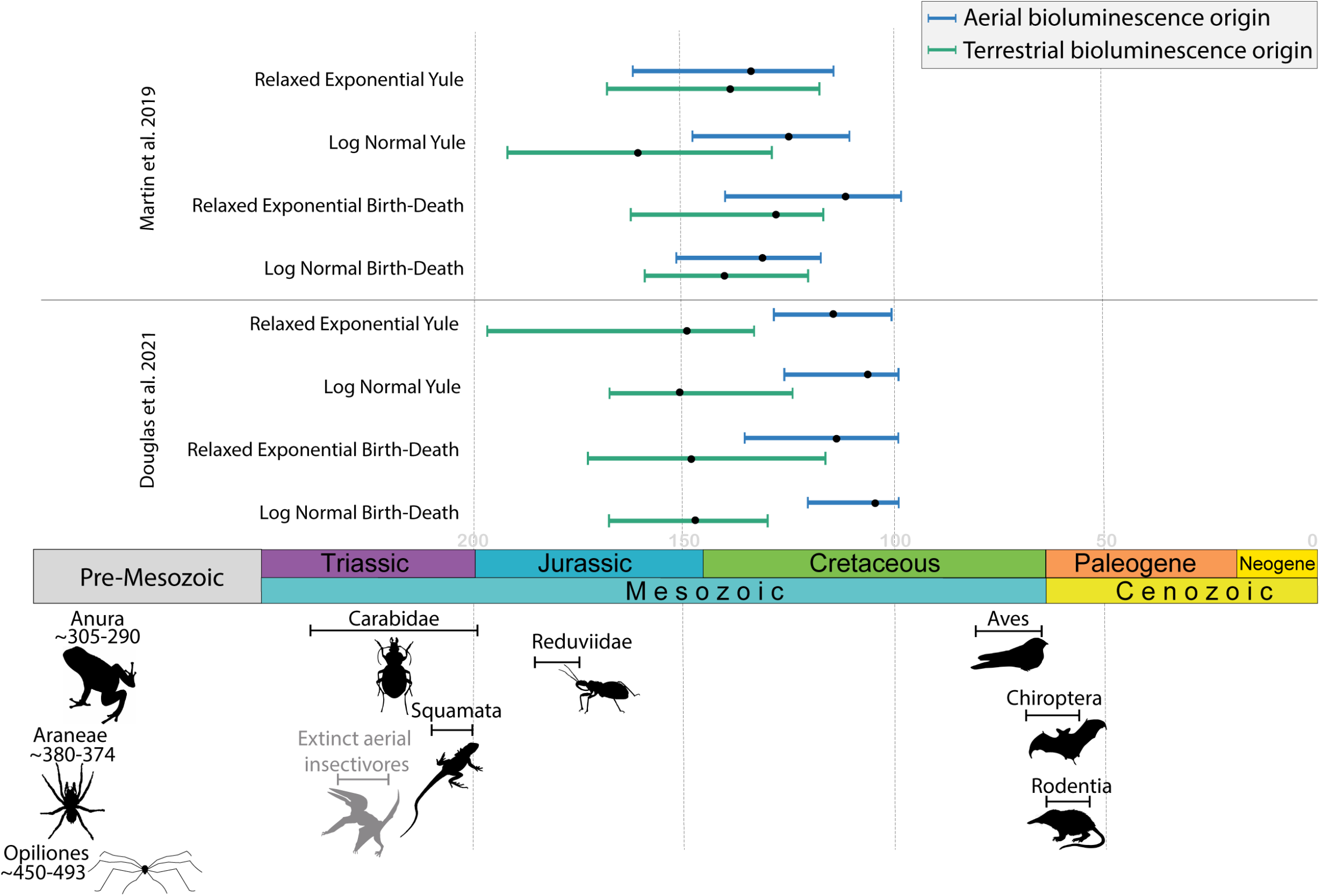
Comparison of divergence time estimates for nodes representing the origins of both terrestrial (blue line) and aerial (blue line) bioluminescence and with published clade origin ages for potential extant (black) and extinct (grey) predator groups. Times are written out for Opiliones, Anura, and Araneae due to age and figure constraints. Dotted vertical lines represent 50 million-year intervals.

### Divergence time estimation

Accurate divergence time estimation relies on a combination of broad taxon sampling, an accurate phylogenetic reconstruction, and sufficient breadth and depth of fossil sampling [44, 45, 75, 76]. A limited fossil record, the difficulty of placing extinct taxa, and the immense extant diversity for a group like Elateroidea (21,000 described species), have led to continued disagreement for the ages of these groups. Previous estimates for the origin of Elateroidea ranged from 220–130 mya, and estimates for the origin of Lampyridae ranged from 130 –75 mya [77, 78, 79]. These large ranges were likely due to a combination of limited taxon sampling for lampyrids and limited fossil calibration points. For example, Bocak et al. [78], hypothesized the age of Elateroidea to be ∼220 mya and Lampyridae as ∼130 mya based on two extant fireflies, 19 total Elateroidea taxa, and a single elateriform fossil calibration point (15 gene dataset). Similarly, McKenna et al. [77] estimated the age of Elateroidea to be ∼130 mya and Lampyridae to be ∼78 mya, with a dataset of four firefly taxa (eight gene dataset). In a subsequent analysis McKenna et al. [77] reconstructed a phylogeny of Coleoptera and estimated the of Elateroidea to be ∼187 mya and ∼120 mya for the branch leading to Lampyridae, however, this study included only a single lampyrid species and no internal elateroid fossil calibration, instead utilizing an ancestral click beetle at the base of Elateroidea+Byrroidea (4800 gene dataset). Oba et al. [28] reconstructed the luciferase gene for the common ancestor of elateroids and estimated the origin for this luciferase at ∼102.55 MY using seven taxa and a single gene tree (18S).

Our analyses based on Martin et al. [40] with 436 loci, 88 firefly species (98 taxa total), and four fossil calibrations across the topology recovered the age of terrestrial bioluminescence as 141.17 (122.63– 161.17) mya and aerial bioluminescence of 133.18 (117.86–152.47) mya. Our analyses based on Douglas et al. [41] with 958 loci, 2 firefly species (an additional 2 “lampyroids”) (88 taxa total), and 7 fossil calibrations. Divergence time estimates for this dataset recovered the root age for Elateroidea as 278.61 (262.03–294.86) mya, the age of terrestrial bioluminescence as 148.03 (130.12–166.80) mya, and the age of aerial bioluminescence at 104.97 (99.00–120.90) mya. It also recovered an independent origin of adult bioluminescence in Elateridae where Pyrophorini branches off at 115.42 (91.70–133.70) mya.

These divergence time estimates for Lampyridae are older than previously published estimates, most likely due to improved taxon sampling. McKenna et al. [77, 80] had limited lampyrid and broader elateroid taxon sampling with limited fossil calibrations that resulted in much younger ages for Lampyridae, and Coleoptera overall, which was already noted by others [45, 81, 82]. Our age estimates are closely aligned to the estimates of Bocak et al. [78], which is likely due to their larger elateroid sampling and ingroup fossil calibrations (e.g., *Elaterophanes*). With updated clade ages, and the species-level biological information gathered for bioluminescence across our phylogeny, we can now examine the validity of previous hypotheses relating to predation and the origin of firefly bioluminescence.

### Age of bioluminescence in beetles

Clade ages for bioluminescence were largely congruent between all different clock and tree models implemented with minor variations in estimated ages reported between each parameter combination (Supp. Table 4). The Yule model does not account for any extinction and elateroid beetles have experienced at least two global extinction events. Thus, we have focused our discussion going forward on the ages resulting from Birth-Death tree models, and those utilizing the Martin et al. [40] dataset as the taxon sampling for bioluminescent taxa is much greater. However, we report the variation in age for each model to demonstrate the robustness of our analyses. The age estimates for each clock model combination largely overlapped for the Birth-Death models; here we discuss the relaxed clock log normal model, which assumes the branch rates are normally distributed and has been shown to more precisely estimate ages and therefore is widely used [83, 84]. Our ancestral state reconstruction placed the origin of terrestrial bioluminescence operating as an aposematic signal at 141.17 (122.63–161.17) mya during the early Cretaceous period. This was followed by the origin of aerial bioluminescence functioning as a sexual signal in extant fireflies, estimated to be at 133.18 (117.86–152.47) mya. Terrestrial bioluminescence is an ancestral trait in Lampyridae and relatives (Phengodidae, and Rhagophthalmidae), preceding the origin of adult aerial bioluminescence that is used widely as a sexual signal in modern fireflies [26, 27, 85]. Assuming that the original function of bioluminescence was predator deterrence, it begs the question as to what predator, or predators, could have driven the origin of firefly bioluminescence.

### Predators

Clade ages for potential predators that fireflies likely encountered during their early history, that would have contributed to the origin of bioluminescence as an aposematic signal, were compiled. We identified potential groups of firefly predators that were hypothesized to feed on these groups and were prevalent and broad enough in distribution to potentially function as significant firefly predators. Terrestrial predators included: Opiliones (harvestmen) [73], Araneae (spiders) [67], Carabidae (carabid beetles) [45], Anura (frog and toads) [70], and Squamata (lizards and snakes) [68], Reduviidae (assassin bugs) [72], Rodentia [71]. Aerial predators included: Aves (birds) [74] and Chiroptera (bats) [69]. An estimation for the origins of terrestrial bioluminescence allows us to discuss the likeliness of these predator groups as drivers of the evolution of warning signals. Several of the terrestrial predator groups, including Anura, Araneae, Carabidae, Opiliones, and Squamata emerged prior to the Jurassic (>200 mya), before the origin of terrestrial beetle bioluminescence and therefore could have contributed to the origin of aposematic beetle bioluminescence that arose 141 mya. Reduviidae emerged around 178 mya; however, most of the diversity in this group did not appear until the Late Cretaceous (∼97 mya) [72].

In contrast, several other insectivore clades (Aves, Chiroptera, Rodentia) originated much later (<70 mya) than beetle bioluminescence and thus could not have been selective agents in either terrestrial or aerial bioluminescence. It has been suggested that modern aerial predators, specifically bats, could have driven the origin of aerial bioluminescence as a predator avoidance strategy [36] This is contradicted by our data, with bats originating at ∼65 mya [69], about 52-87 million years after the origin of aerial bioluminescence in fireflies. This result supports the idea that the original purpose of adult, aerial bioluminesence was that of sexual signaling. However, it is possible that once bioluminescence was used by adult fireflies in aerial displays, potential aerial predators like bats and birds could utilize adult firefly bioluminescence as an additional cue to avoid distasteful prey [36]. The sensory systems of insectivorous bats can detect both the light spectrum of firefly light emissions and the ultrasonic wingbeat clicks emitted by flying fireflies [86, 87]. In addition, bats can learn to discriminate between flying insects based on their different echo signatures in their echolocation calls [88, 89] and they make adaptive prey selection decisions to increase profitability [90]. Bats have indeed been shown to reject flying fireflies based on their chemical defenses, reinforced by their sonar profile and/or bioluminescent signal that speed up avoidance learning [36]. Whether predators like bats impose selection on the aerial bioluminescence of fireflies and possibly contribute to the maintenance of aerial light signals in beetles remains to be tested. Other predators such as nocturnal birds and rodents [37] that could have preyed on bioluminescent fireflies also originated significantly after the origin of both terrestrial and aerial bioluminescence (Fig. 2). For example, modern birds originated ∼75 mya [74] and rodents at ∼61 mya [71], placing them ∼65-79 million years after the estimated origin of terrestrial bioluminescence and ∼58-72 million years after aerial bioluminescence (Fig. 2). Our age estimates for bats, birds, and rodents are almost certainly older than the actual insectivorous lineages within each. Given that beetle larval bioluminescence evolved first and operated in a terrestrial environment, aerial predators, such as bats, could not have been the original receivers or drivers of these aposematic signals. The signal would have been directed toward contemporary predators of elateroids in the early Cretaceous.

There are several extinct vertebrate insectivore groups that could have also preyed on lampyrids and other elateroid beetles prior to or during the early Cretaceous (>100 mya) [91, 92]. Most of the early mammalian insectivores are more limited in both known diversity and distribution in the fossil record [92] and would thus likely have only added to an already existing predation pressure in the terrestrial environment. One potential exception could have been the aerial pterosaurs [93]. Pterosaurs were the first vertebrate group to develop true flight in the late Triassic (∼229 mya), with the common ancestor of the group hypothesized to be insectivorous [94]. Furthermore, Pterosauria had a broad enough distribution to have encountered bioluminescent beetles [95], and some species are assumed to have been crepuscular or even nocturnal [96], and thus could have been early receivers of the aposematic signals of fireflies and other elateroid beetles. Although tantalizing, there is limited evidence to support that pterosaurs fed on elateroid beetles.

In summary, our extensive taxon sampling and numerous well-placed fossils across each topology, recovered well-supported and congruent ages for bioluminescent beetles. These ages are older than previous estimations and preclude modern aerial predators as selective agents for the origin of beetle bioluminescence. However, several groups of modern terrestrial predators predate the origin of bioluminescence and thus could have functioned as selective agents for the origin of terrestrial bioluminescence in fireflies, and possibly other elateroid beetles. Based on the present day use and abundance of terrestrial bioluminescence as aposematic signals and the presence of terrestrial insectivores at the origin of beetle bioluminescence strongly suggests that terrestrial beetle bioluminescence arose as an aposematic signal. In contrast, aerial beetle bioluminescence, which is widely used as a sexual signal during mate search in extant beetles, likely originated as such. We do not rule out that extant insectivores such as bats, birds, and rodents, that emerged in the fossil record after the origin of firefly bioluminescence, may still operate as contributing factors in maintaining bioluminescence. However, it is now clear that the origin of aerial bioluminescence in fireflies predates any extant aerial predator group. We find that adult firefly bioluminescence outshines, or predates, the origins of extant aerial predators by roughly 60 million years.

## Acknowledgments

We dedicate this work to the late Dr. Jim Lloyd, *the* giant of firefly research who mentored us all in significant ways, oftentimes with humor and colorful language. We thank all those that provided specimens or identifications for the original phylogenetic efforts. We would also like to thank the reviewers and editorial staff for suggestions that greatly improved the manuscript. L.B. and D.K. were partly funded by GACR 22-33714S. This research was in part funded by the National Science Foundation (DEB-1655981 S.M.B; DEB-1655908 K.S.H; and DEB-1655936 M.A.B).

