## Supplemental Table 1 for "Beetle bioluminescence outshines aerial predators"

**Supplementary Material**

| **Fossil** | **Age (mya)** | **Clade** | **Dataset** |
|---|---|---|---|
| *Phausis fossilis* | 25 | Lamprohizinae | Martin |
| *Photinus kazansevi* | 36.2 | *Photinus* | Martin |
| *Cardiophorus exhumatus* | 48 | *Cardiophorus* | Douglas |
| *Lissomus taxodii* | 59.2 | Lissominae | Douglas |
| *Cretophengodes azari* | 99 | (Rhagophthalmidae + Phengodidae) | Both |
| *Protoluciola albertalleni* | 99 | Luciolinae | Both |
| *Ganestrius elongatus* | 148 | Negastriinae | Douglas |
| *Ageratus delicatus* | 149 | Agrypninae | Douglas |
| *Elateridium* spp. | 242 | Elateridae | Douglas |

Supp. Table 1: Fossils used in divergence time estimation with their relative age, placement, and dataset they were applied to.
