## Supplemental Table 3 for "Beetle bioluminescence outshines aerial predators"

| Predator Group | Estimated Origin (Ma) | Reference |
|---|---|---|
| Opiliones | 450–493 | [73] |
| Araneae | 374–380 | [67] |
| Extinct aerial predators | 234–224 | [97] |
| Squamata | 200–210 | [68] |
| Carabidae | 200–240 | [45] |
| Anura | 290–305 | [70] |
| Reduviidae | 176-185 | [72] |
| Aves | 67–82 | [74] |
| Chiroptera | 58–71 | [69] |
| Rodents | 56–66 | [71] |

Supp. Table 3. Estimated ages of each firefly predator group compiled from the literature.
