## Supplemental Table 4 for "Beetle bioluminescence outshines aerial predators"

| Tree | Clock | Terrestrial | Aerial | Dataset |
|---|---|---|---|---|
| BD | RCLN | 141.17 (122.63–161.17) | 133.18 (117.86–152.47) | Martin |
| BD | RCE | 129.87 (117.39–164.44) | 110.30 (99.36–140.97) | Martin |
| Yule | RCLN | 160.77 (129.73–192.69) | 124.06 (112.65–148.25) | Martin |
| Yule | RCE | 139.51 (118.78–169.69) | 134.60 (115.69–163.76) | Martin |
| BD | RCLN | 148.03 (130.12–166.80) | 104.97 (99.00–120.90) | Douglas |
| BD | RCE | 147.92 (116.00–174.12) | 113.12 (99.00–137.94) | Douglas |
| Yule | RCLN | 150.42 (123.31–166.68) | 108.72 (99.00–125.39) | Douglas |
| Yule | RCE | 146.91 (135.40–195.35) | 117.49 (107.37–129.21) | Douglas |

Supp. Table 4. Divergence time estimates for nodes reconstructed to be the origins of terrestrial and aerial bioluminescence under different tree and clock models with each dataset. RCLN- Relaxed Clock Log Normal, RCE- Relaxed Clock Exponential, BD- Birth-Death.
