## Supplemental Figure 1 for "Beetle bioluminescence outshines aerial predators"

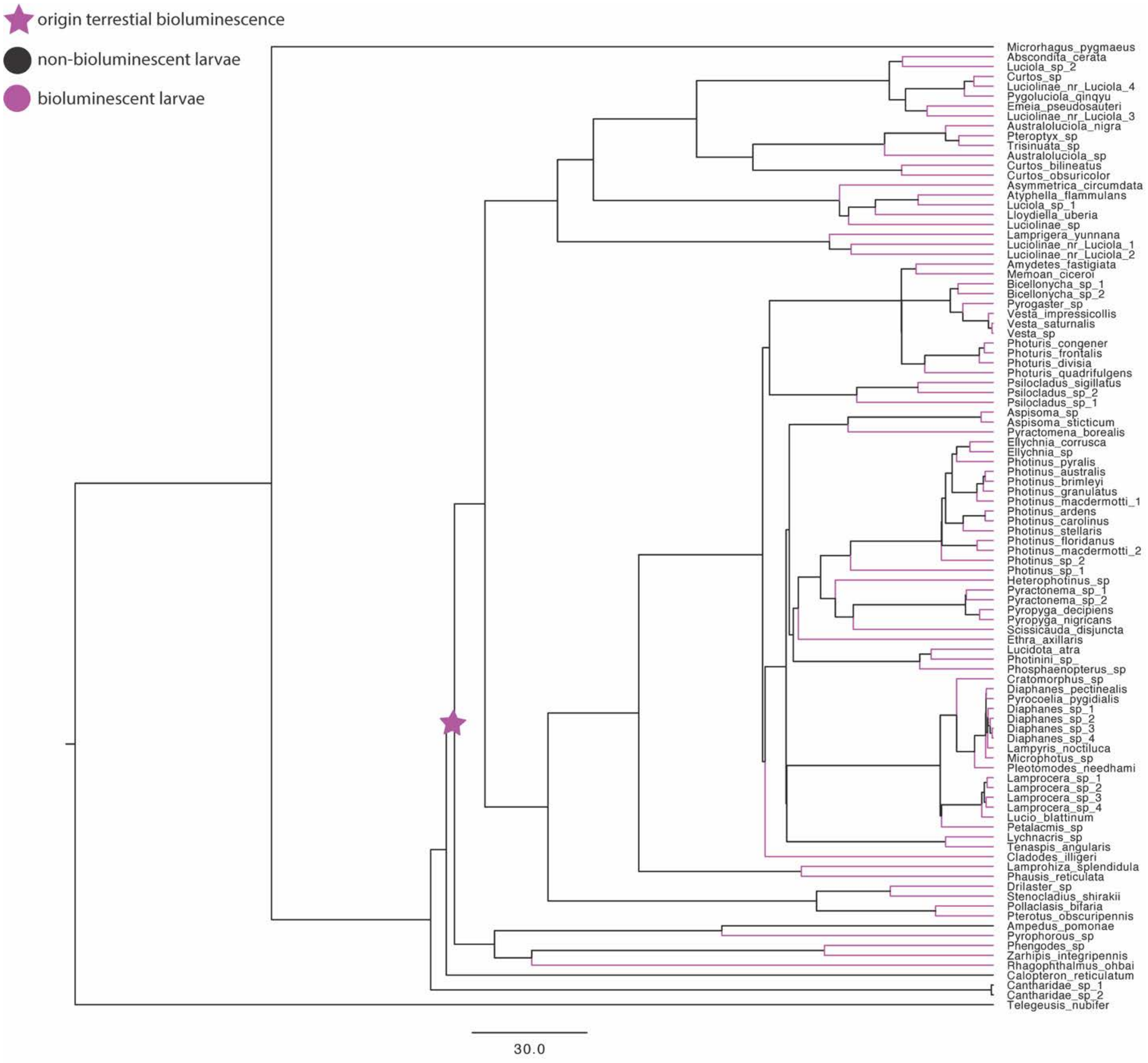


Supp. Figure 1. Maximum parsimony ancestral state reconstruction of Martin et al. [40] phylogeny for larval bioluminescence.

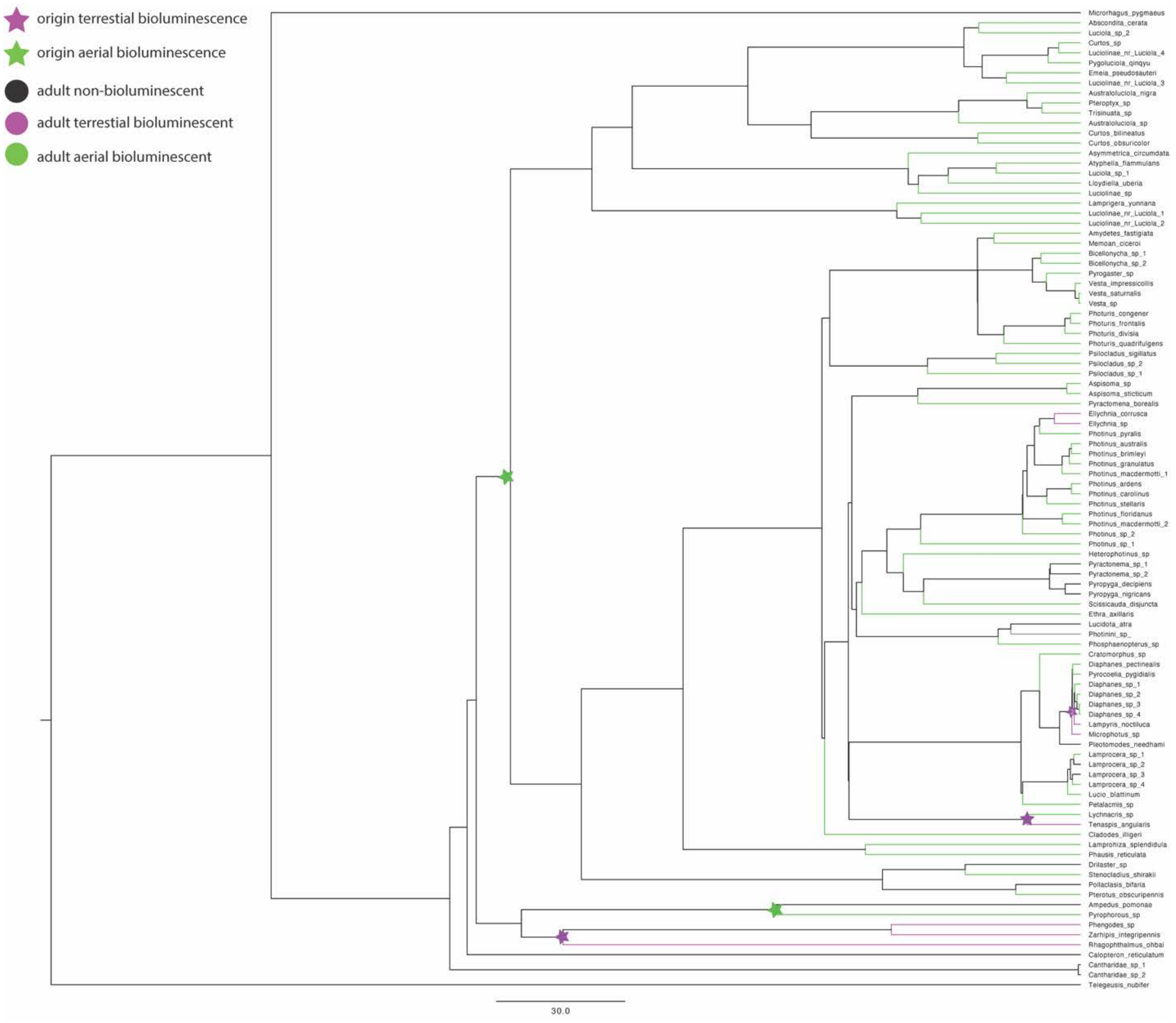


Supp. Figure 2. Maximum parsimony ancestral state reconstruction of Martin et al. [40] phylogeny for adult bioluminescence. Star marked with “2” signifies two independent origins of terrestrial bioluminescence. Gray branches indicate the signal state is unknown.

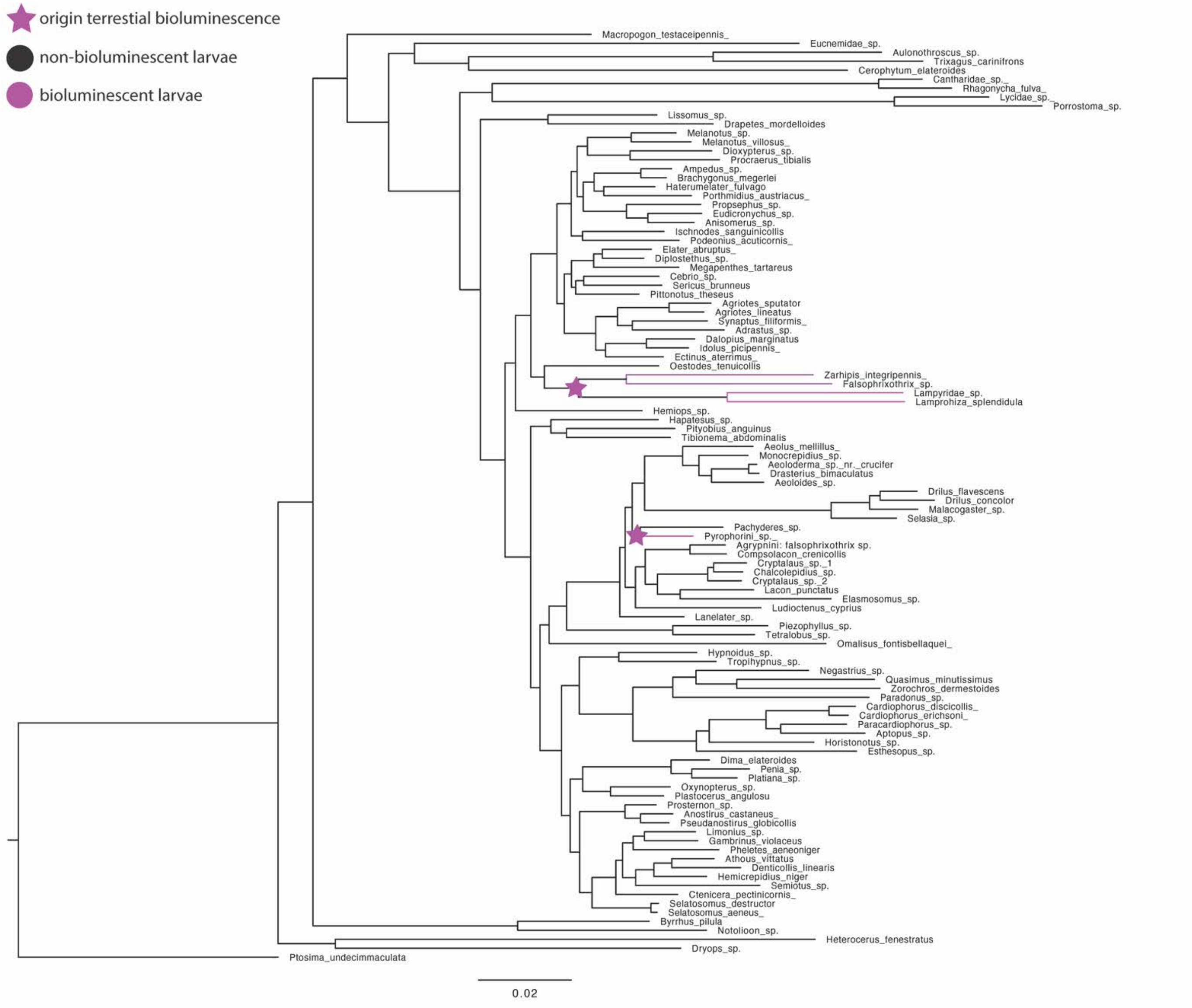


Supp. Figure 3. Maximum parsimony ancestral state reconstruction of Douglas et al. [41] phylogeny for larval bioluminescence.

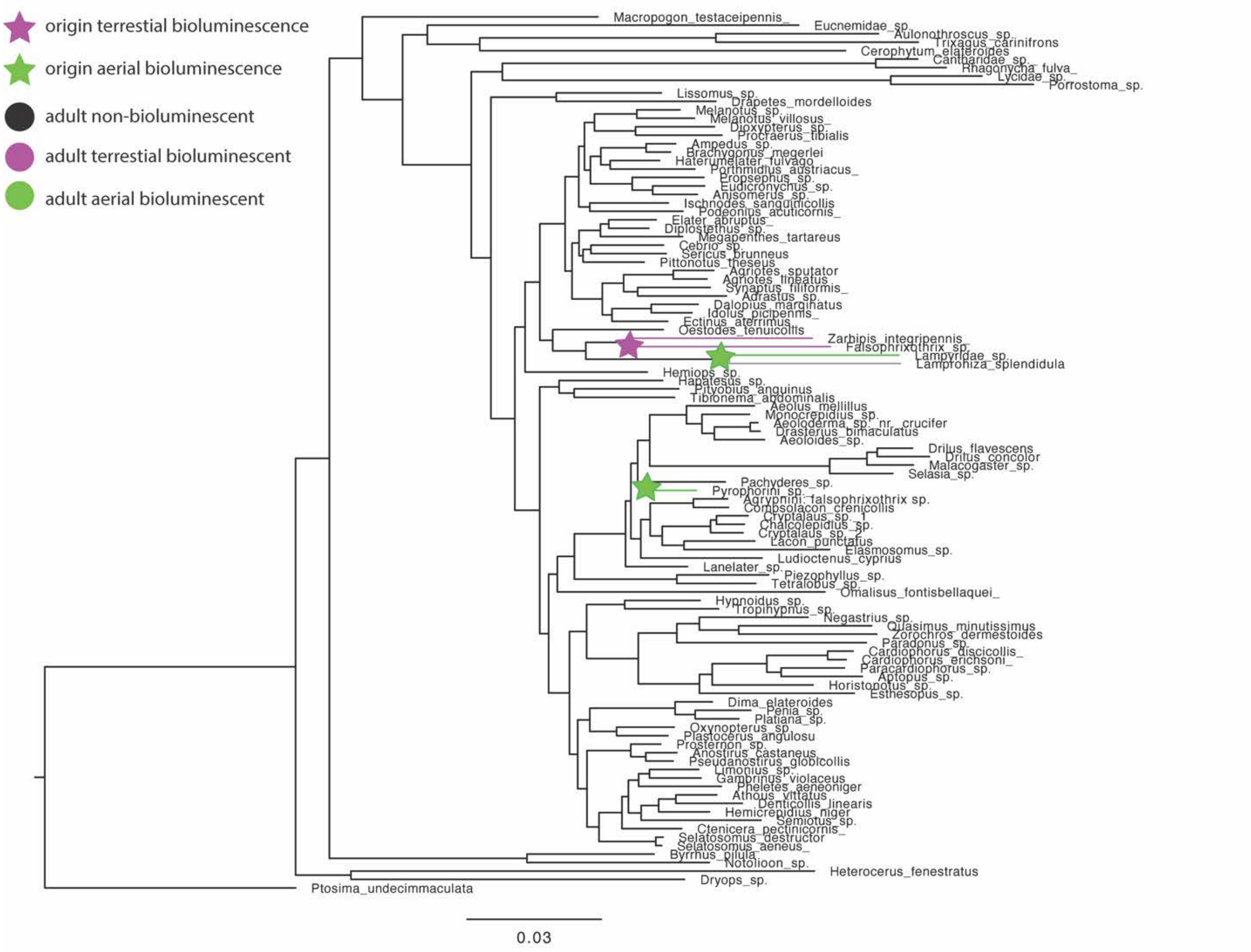


Supp. Figure 4. Maximum parsimony ancestral state reconstruction of Douglas et al. [41] phylogeny for adult bioluminescence. Gray branches indicate the signal state is unknown.
