## Supplemental Figure 2 for "Beetle bioluminescence outshines aerial predators"

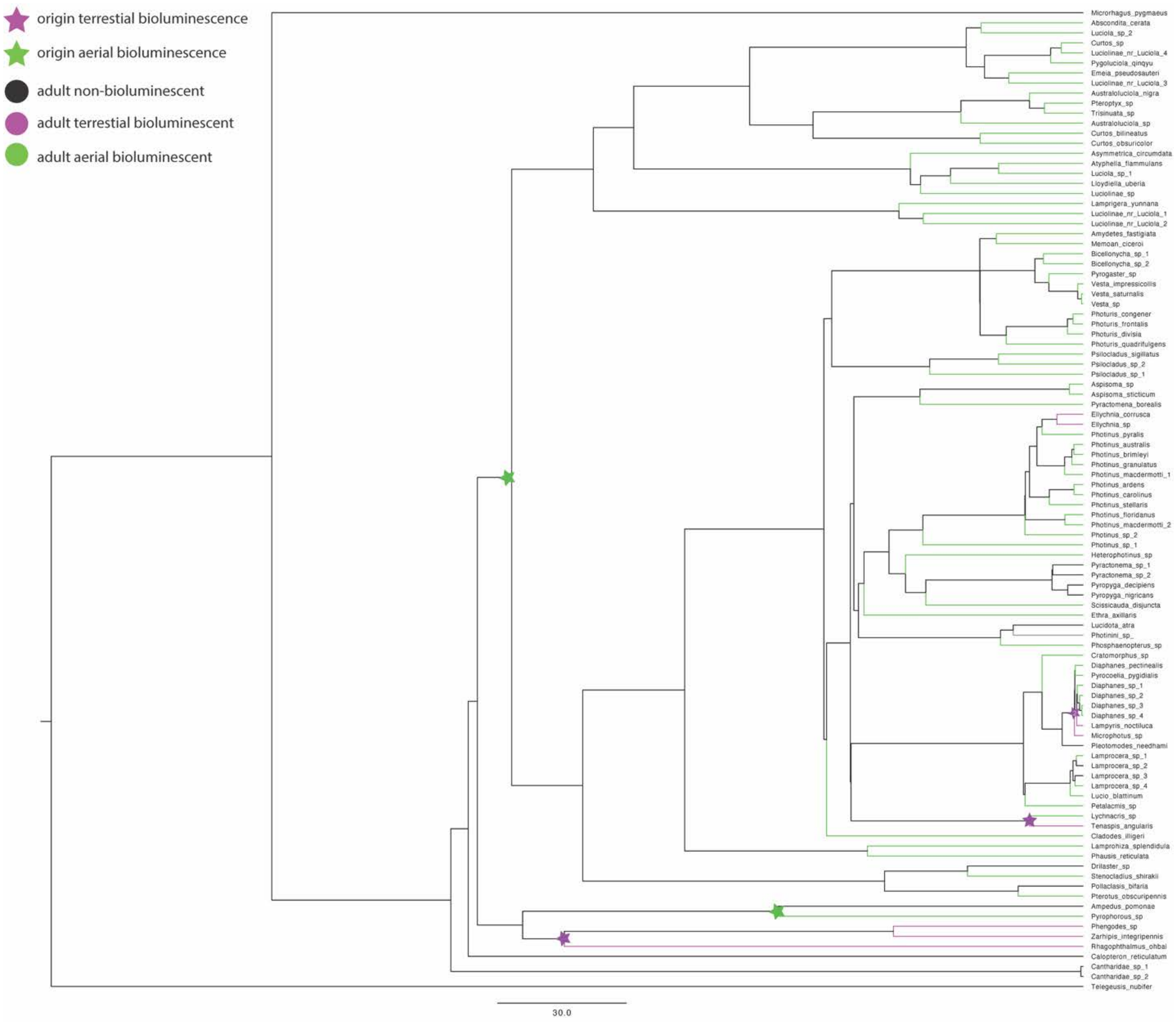


Supp. Figure 2. Maximum parsimony ancestral state reconstruction of Martin et al. [40] phylogeny for adult bioluminescence. Star marked with “2” signifies two independent origins of terrestrial bioluminescence. Gray branches indicate the signal state is unknown.
