## Supplemental Figure 3 for "Beetle bioluminescence outshines aerial predators"

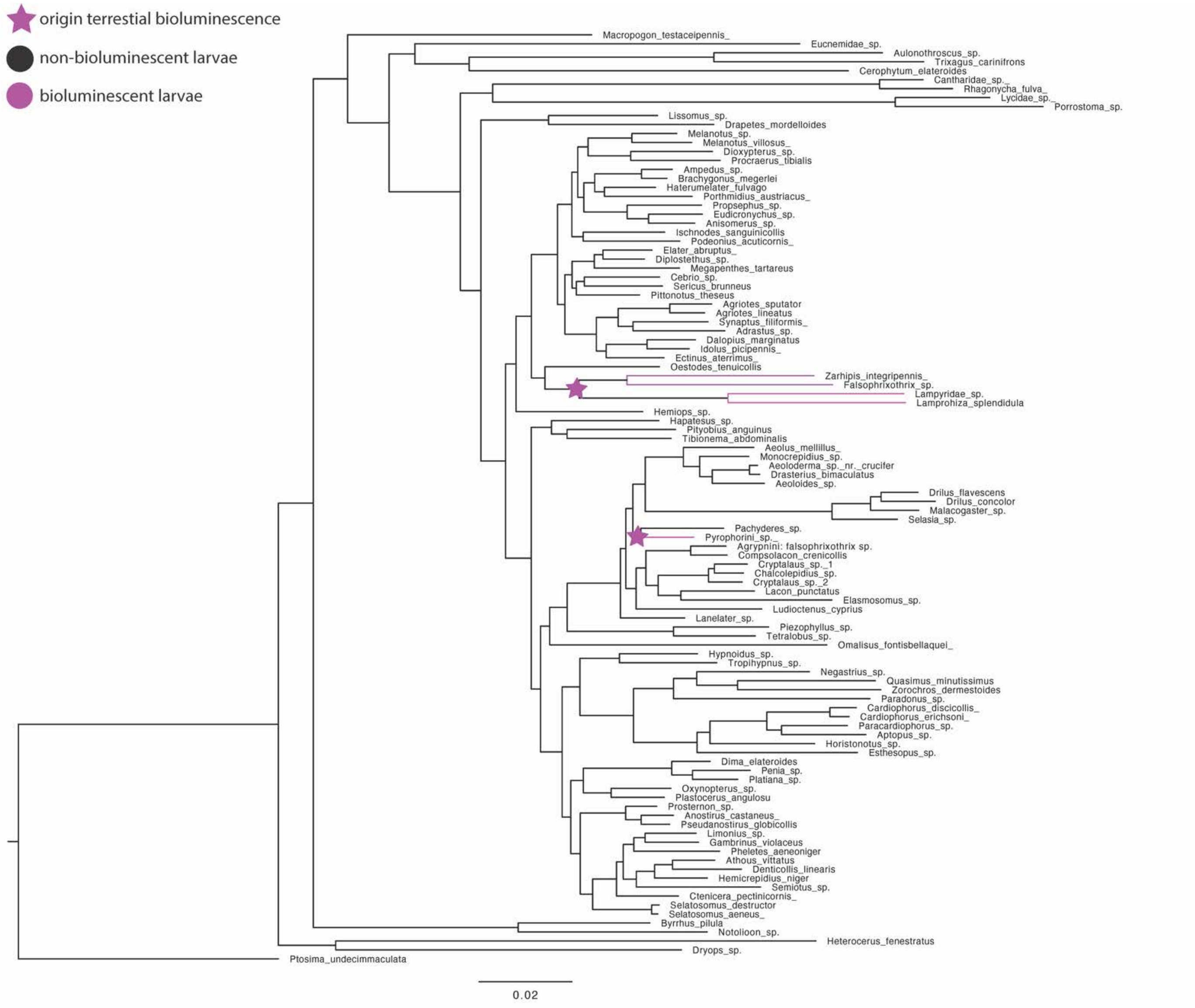


Supp. Figure 3. Maximum parsimony ancestral state reconstruction of Douglas et al. [41] phylogeny for larval bioluminescence.
