## Supplemental Figure 4 for "Beetle bioluminescence outshines aerial predators"

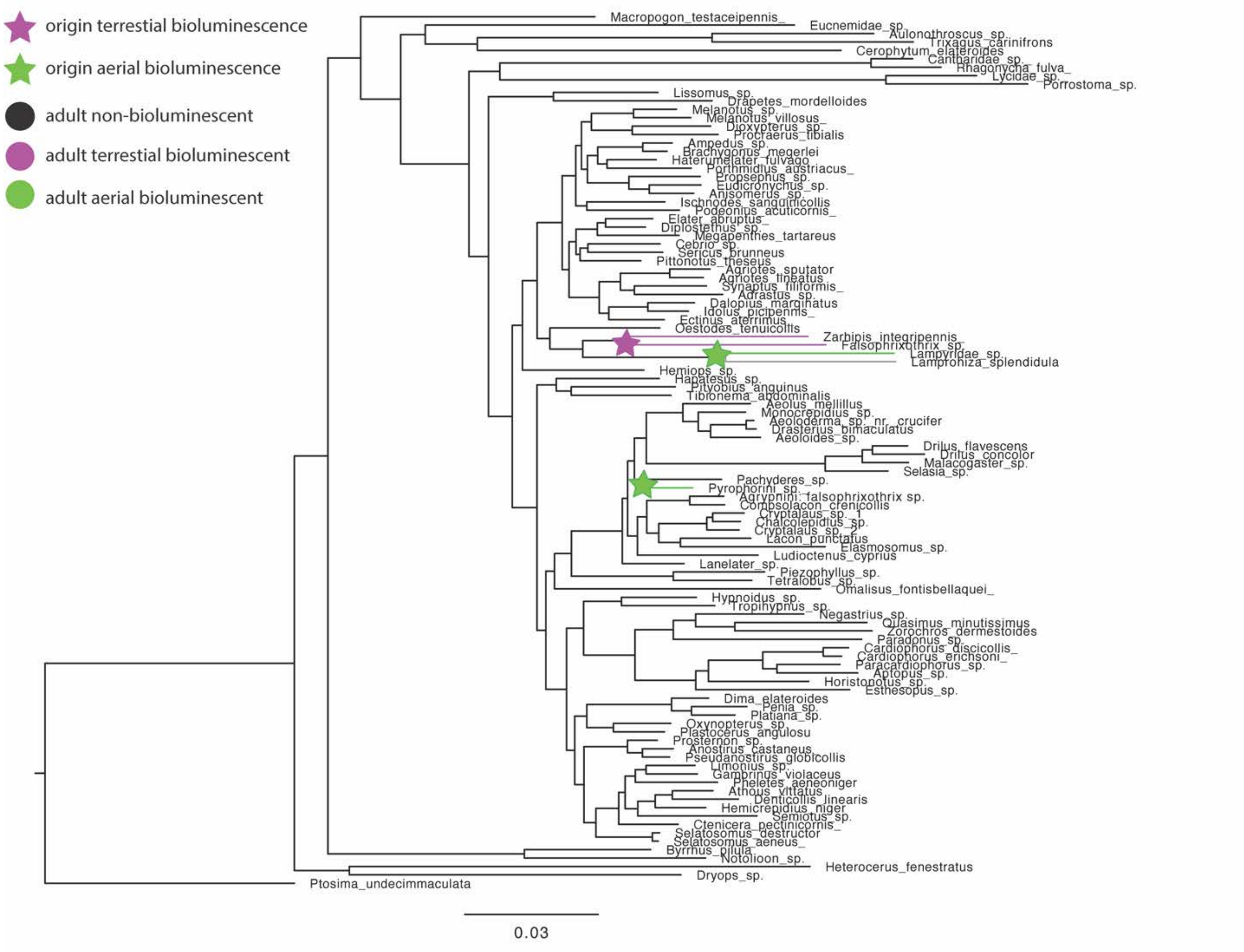


Supp. Figure 4. Maximum parsimony ancestral state reconstruction of Douglas et al. [41] phylogeny for adult bioluminescence. Gray branches indicate the signal state is unknown.
